# Core Fermentation (CoFe) granules focus coordinated glycolytic mRNA localization and translation to fuel glucose fermentation

**DOI:** 10.1101/741231

**Authors:** Fabian Morales-Polanco, Christian Bates, Jennifer Lui, Joseph Casson, Clara A. Solari, Mariavittoria Pizzinga, Gabriela Forte, Claire Griffin, Harriet E. Burt, Hannah L. Dixon, Simon Hubbard, Paula Portela, Mark P. Ashe

**Affiliations:** School of Biological Sciences, Faculty of Biology, Medicine and Health, The University of Manchester, Manchester Academic Health Science Centre, Michael Smith Building, Oxford Rd., Manchester. M13 9PT. UK; Department of Biological Sciences, Stanford University, 318 Campus Drive, Stanford, CA 94305; Departamento de Química Biológica, Facultad de Ciencias Exactas y Naturales, Universidad de Buenos Aires, Buenos Aires, Argentina. IQUIBICEN-CONICET; MRC Toxicology Unit, University of Cambridge, Hodgkin Building, Lancaster Road, Leicester. LE1 9HN. UK

**Author notes:** These authors contributed equally to this paper (Fabian Morales-Polanco, Christian Bates).

**Keywords:** Glycolysis, mRNA translation, phase separation, RNA granules, translation factories

## Abstract

Glycolysis is a fundamental metabolic pathway for glucose catabolism across biology, and glycolytic enzymes are amongst the most abundant proteins in cells. Their expression at such levels provides a particular challenge. Here we demonstrate that the glycolytic mRNAs are localized to granules in yeast and human cells. Detailed live cell and smFISH studies in yeast show that the mRNAs are actively translated in granules, and this translation appears critical for the localization. Furthermore, this arrangement is likely to facilitate the higher level organisation and control of the glycolytic pathway. Indeed, the degree of fermentation required by cells is intrinsically connected to the extent of mRNA localization to granules. On this basis, we term these granules, Core Fermentation (CoFe) granules; they appear to represent translation factories allowing high-level co-ordinated enzyme synthesis for a critical metabolic pathway.

## Introduction

The glycolytic pathway lies at the core of metabolic activity as a virtually ubiquitous biochemical pathway across living cells. The pathway serves both to supply energy and maintain levels of biochemical intermediates (Bar-Even et al., 2012). Multiple genes express a variety of isoforms for many glycolytic enzymes providing abundant scope for adaptable regulation (Masters et al., 1987; Oparina et al., 2013; Postmus et al., 2012; Warmoes and Locasale, 2014). The pathway was gradually pieced together by a succession of influential biochemists including Meyerhof, Embden, and Parnas (Bar-Even et al., 2012; Barnett, 2005; Schurr and Gozal, 2015). After these major biochemical breakthroughs, interest in central metabolism waned over a period where it was often perceived to perform mundane ‘housekeeping’ functions (Bar-Even et al., 2012; Ray, 2010). More recently, the pathway and its regulation have received renewed interest for various reasons: including connections to cancer and cellular proliferation (Diaz-Ruiz et al., 2011; Gill et al., 2016), moonlighting activities of the glycolytic enzymes (Castello et al., 2015; Kim and Dang, 2005) and increased interest in central metabolism as a focus for metabolic engineering in a synthetic biology era (Lim and Jung, 2017).

In many aerobic cells, the pyruvate produced by glycolysis is transported to and oxidised in the mitochondria via respiration (Gray et al., 2014). However, under anaerobic conditions and in various aerobic cells, such as yeast, lymphocytes and cancer cells, glucose is fermented through to ethanol or lactic acid. Hence, in these cells glycolysis serves as the major source of ATP and intermediates (Lunt and Vander Heiden, 2011). Indeed, the reduction of pyruvate to ethanol or lactic acid can be viewed as an extension of glycolysis.

Given the critical nature of the glycolytic pathway in energy production and cellular metabolism, it is unsurprising to find that the pathway is regulated by a myriad of different mechanisms. These include direct regulation of the enzymes via substrate and product concentration (Wegner et al., 2015), allosteric enzyme regulation by small molecules (Shen et al., 2016) and post-translational covalent modifications (Shenton and Grant, 2003; Tripodi et al., 2015). Aside from controls of enzymatic activity, other regulatory mechanisms act at the level of gene transcription (Chambers et al., 1995; Yeung et al., 2008), mRNA processing/ stability (Krieger and Ernst, 1994; Lunghi et al., 2015), protein stability (Benanti et al., 2007; Lu et al., 2014; Riera et al., 2003), translation (Daran-Lapujade et al., 2007; Man and Pilpel, 2007) and protein localization (Jin et al., 2017). While clearly the glycolytic enzymes and the mRNAs that encode them *can* be regulated, they are often viewed as providing ‘housekeeping’ functions. Indeed in yeast, many of the glycolytic mRNAs are amongst the most abundant, heavily translated mRNAs in the cell. This raises obvious questions, such as how is this level of gene expression attained both at the transcriptional and post-transcriptional levels? Furthermore, how is this scale of gene expression co-ordinated across the pathway such that appropriate levels of enzyme are produced to generate a metabolic flux that is pertinent to the cellular conditions?

A number of recent observations have supplemented understanding of glycolysis and the role of glycolytic enzymes in cells. For instance, it has become evident that a number of glycolytic enzymes ‘moonlight’ as RNA binding proteins (Castello et al., 2015). Indeed it has been suggested that many of the glycolytic proteins bind to glycolytic mRNAs to orchestrate control of the pathway (Matia-Gonzalez et al., 2015). In addition, the localization of two glycolytic mRNAs in yeast, *PDC1* and *ENO2*, has been identified as important in their translation control and in the formation of mRNA processing bodies or P-bodies (PBs) after glucose starvation (Lui et al., 2014).

mRNA localization has been commonly considered as a means to generate localized sources of protein, with specific examples involved in cellular polarization identified across many biological systems-*ASH1* mRNA in yeast (Long et al., 1997), *bicoid* in *Drosphila* oocytes (Berleth et al., 1988) and *Vg1* in *Xenopus* oocytes (Melton, 1987). In these cases, translationally repressed mRNAs are localised in a transit process involving motor proteins and cytoskeletal elements (Besse and Ephrussi, 2008). Another situation where translationally repressed mRNAs become localized is under stress conditions, where non -translated mRNAs can enter either PBs or stress granules (SGs) to play roles in mRNA degradation and/or storage (Hoyle and Ashe, 2008; Hubstenberger et al., 2017; Jain and Parker, 2013). More global assessments of mRNA localization suggest that the phenomena is widespread: large numbers of mRNA species are localized in *Drosophila*, neuronal cells and yeast (Gadir et al., 2011; Lecuyer et al., 2007; Miyashiro et al., 1994; Pizzinga and Ashe, 2014; Zipor et al., 2009; Zivraj et al., 2010). Even so, mRNA localization is rarely thought to play a role in core housekeeping functions such as central metabolism.

In this study, we show that glycolytic mRNAs in yeast and human cells are specifically localized to granules. In yeast, we define the <u>Co</u>re <u>Fe</u>rmentation (CoFe) granule, a core glycolytic mRNA granule where glycolytic mRNAs are colocalized and cotranslated. Translation is a prerequisite for CoFe granules and individual mRNA translation is required for localization. Finally, we show that the presence of mRNA granules correlates with the degree of glycolytic function required by the cell. We suggest that the localization of these mRNAs provides a means to generate the scale of protein expression required for such a critical pathway and permit rapid co-ordinated regulation.

## Results

### Glycolytic mRNAs localize to granules under active growth conditions

Previous work from our laboratory has highlighted that the glycolytic mRNAs, *PDC1* and *ENO2*, encoding pyruvate decarboxylase and enolase, respectively, are translated in cytoplasmic granules (Lui et al., 2014) (Figure 1A). To evaluate whether glycolytic mRNAs in general are localized these sites, we have utilized the m-TAG system, where elements of the MS2 bacteriophage are used to tether GFP to an mRNA to study its localization in live cells (Haim-Vilmovsky et al., 2011). Accordingly, MS2 stem loops were directly inserted into the 3’UTR sequences of the glycolytic genes at their endogenous genomic loci. The localization of the resulting mRNAs was then followed via co-expression of the MS2 coat protein-GFP fusion (MS2-CP-GFP). It should be noted that MS2-CP-GFP expression alone generates diffuse fluorescence throughout the cell (Figure 1A). In contrast, when MS2 stem loops are integrated into glycolytic mRNA 3’UTRs, the vast majority of the resulting mRNAs are observed in granules (Figure 1B). This includes mRNAs that encode enzymes at every step of the glycolytic pathway (Figure 1C). Notably, not all MS2-tagged mRNAs localize to granules of this kind (Lui et al., 2014; Simpson et al., 2014), for instance, two non-glycolytic mRNAs, *GLO1* and *PFK26*, where the gene products are involved in the control of glycolysis, and the glycolytic mRNA, *PYK2*, are not observed in granules (Figure 1B). Previously *PDC1* and *ENO2* mRNAs were shown to localize to ~20 granules per cell (Lui et al., 2014), and here we show that 14 of the 15 tested glycolytic mRNAs localize to a similar number of granules (Figure 1D).

**Figure 1.**
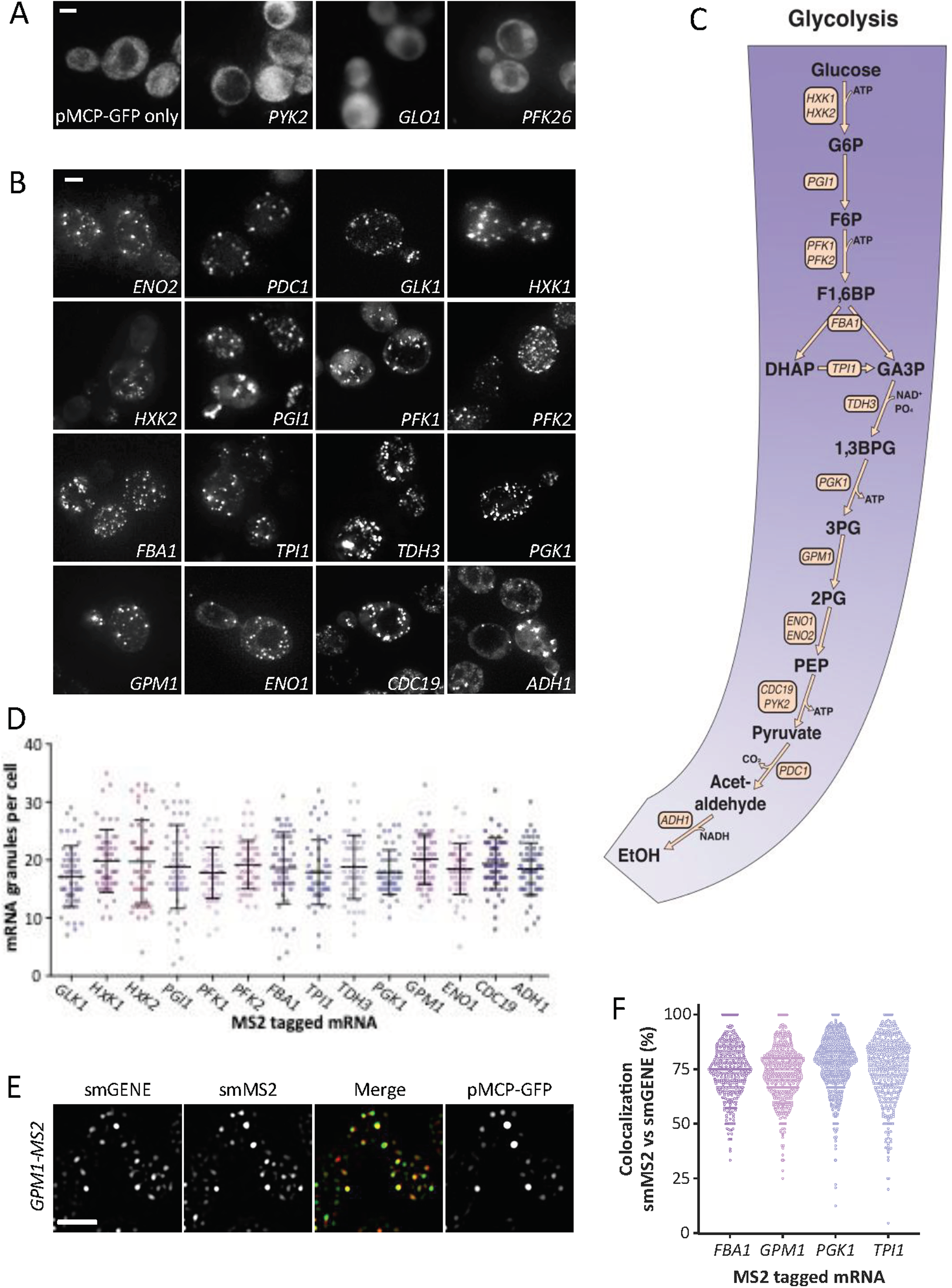
MS2-tagged glycolytic mRNAs are localized to granules in *S. cerevisiae*. (A) and (B), *z*-stacked images of strains expressing *MS2*-tagged mRNAs as labelled and the MS2 coat protein GFP fusion. Scale bar: 2μm. (C) diagram of glucose fermentation to ethanol depicting the glycolytic mRNAs investigated in this study. (D) A dotplot showing the variation in the number of granules per cell for each of the *MS2*-tagged strains above. n=50. The mean ± SD are indicated for each strain. (E) *z*-stacked images of smFISH performed on strains expressing *MS2*-tagged mRNAs and the MS2 coat protein GFP fusion. smFISH was performed for the canonical *GPM1* gene (smGENE) or the *MS2* stem loop sequence (smMS2). Scale bar: 3μm. (F) Beeswarm plot showing the proportion of smMS2 foci that colocalize with smGENE foci for a subset of strains expressing *MS2*-tagged glycolytic mRNAs. Each dot represents a single cell. n>300.

Recent commentaries have highlighted the potential for an accumulation of fragments of mRNA carrying MS2 stem loops that can impact upon the interpretation of experiments using MS2 tethering systems (Garcia and Parker, 2015, 2016; Haimovich et al., 2016; Heinrich et al., 2017). It is possible that such fragments would accumulate at sites of mRNA degradation. To assess whether this is the case for the granules observed here, a range of approaches were taken. Firstly, it should be noted that all of the experiments presented are conducted on cells actively growing in nutrient replete media. Under these conditions in our experiments, PBs are largely absent (Lui et al., 2014), so the high level accumulation of mRNA fragments at sites of mRNA decay seems unlikely. Secondly, while most of the glycolytic mRNAs tested are present in granules, the MS2 stem loops have a highly variable impact on the steady state level of the mRNAs: some mRNAs are stabilised, others destabilised and some remain unchanged (Figure S1A). This profile is not consistent with MS2 fragments explaining the observed localization. Thirdly, the major mRNA species observed on Northern blots under active growth conditions for either the non-tagged or MS2-tagged *ENO2* and *PDC1* mRNAs were full -length mRNAs (Figure S1B). In contrast, in stressed cells where PBs are present, such as shortly after glucose depletion, degradation fragments for the MS2-tagged mRNAs comprise a high proportion of the total mRNA (Figure S1B). Fourthly, a subset of granule localising glycolytic mRNAs have been tagged with a version of the MS2 system (termed the ‘version 6’ system) where MS2 fragments do not accumulate (Tutucci et al., 2018). This new MS2 system reveals an identical granular pattern of glycolytic mRNA localization to the original MS2 system (Figure S1C). Finally, a single molecule fluorescent in situ hybrdization (smFISH) strategy where probes were targeted to either the MS2 stem loops or the body of the mRNA revealed greater than 75% signal overlap between these two probes (Figure 1E and F). This result suggests that the MS2 region of the mRNA reports the localization of full-length mRNAs. In addition, significant overlap is seen between the MS2-CP-GFP protein signal and either the MS2 RNA probe signal or mRNA body probe signal suggesting that the GFP signal also reports full length mRNAs(Figure 1F). This overlap is observed despite the fact that the MS2-CP-GFP signal is only seen for those granules that exceed a specific intensity threshold, since as reported previously (Pizzinga et al., 2019) the MS2 live cell system only detects multi -mRNA granules. Overall, the combination of different validatory analyses used show that glycolytic mRNAs evaluated using the m-TAG system localize to multi-mRNA granules.

However, insertion of MS2 stem loops could still alter some aspect of an MS2 -tagged mRNAs fate. Therefore, in order to provide an independent assessment of the glycolytic mRNA localization, endogenous unmodified mRNAs were evaluated using smFISH. smFISH strategies commonly use ~30-50 fluorescently labelled oligonucleotides which are hybridized to mRNAs in fixed cells (Pizzinga et al., 2019; Tsanov et al., 2016). Since many glycolytic genes are present in yeast as multiple paralogues with very high levels of sequence identity, the use of smFISH to unambiguously study the localization of individual mRNA species is problematic. Therefore, sets of smFISH probes were designed to study the localization of mRNAs encoded by glycolytic genes that either lack paralogues or harbor substantial sequence differences to their paralogues. As a result, four different glycolytic mRNAs, *GPM1*, *FBA1*, *TPI1* and *PGK1* were analyzed and shown to localize specifically to granules (Figure 2A). In terms of the number of granules per cell, the smFISH data for endogenous mRNAs are entirely complementary to the live cell MS2-tagged mRNAs (*cf*. Figure 2B and 1D). One of the key advantages of the smFISH versus the m-TAG experiments is that for the smFISH data, single mRNA foci are visible. This allows an estimate of mRNA copy number per cell and the proportion of single mRNAs present in multi-mRNA granules (Pizzinga et al., 2019). From the data, it is clear that ~70% of the glycolytic mRNA molecules are present in large granules (Figure 2C). The results correlate well with live cell m-TAG data, where a similar fraction of the *PDC1* and *ENO2* mRNAs were previously found in multi-mRNA granules (Lui et al., 2014). For the non-glycolytic *NPC2* mRNA, which is not localized to large multi-mRNA granules (Pizzinga et al., 2019), a speckled pattern is observed with a reduced, homogeneous fluorescent intensity for each speckle relative to the glycolytic mRNAs. The quantitation of fluorescent intensity profiles shows that the *NPC2* mRNA is rarely localized to multi-mRNA granules and even where *NPC2* multi-mRNA granules can be identified, the proportion of mRNA present is very low (Figures 2A-C). Overall these data confirm that glycolytic mRNAs are housed in large cytoplasmic bodies or granules and, combined with our concurrent studies on translation factor mRNAs (Pizzinga et al., 2019), indicate that the localization observed with the MS2 system in actively growing cells can be representative of and meaningful to the localization observed for an untagged mRNA.

**Figure 2.**
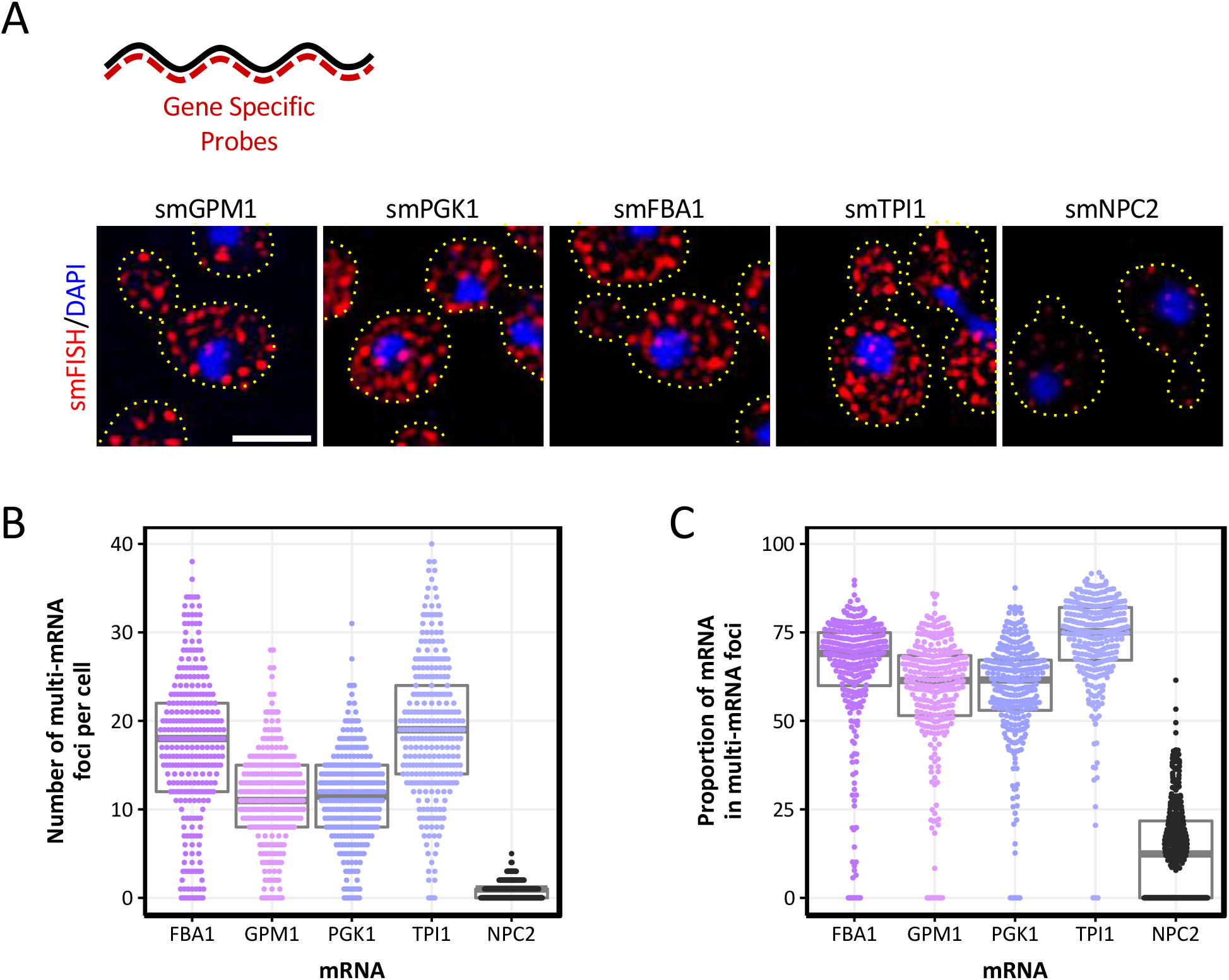
smFISH analysis reveals that endogenous glycolytic mRNAs are present in multi-mRNA granules. (A) Upper diagram depicts the smFISH strategy. Multiple probes complementary to an mRNA (black) are tagged with a specific fluorophore (red). Lower panels show *z*-stacked smFISH images performed for the indicated endogenous glycolytic mRNAs. Both mRNAs (red) and nuclei (blue) are shown. Dotted lines represent the extent of the cell from brightfield micrographs. Scale bar: 3μm. (B) Beeswarm plot showing the number of multi-mRNA (>2.5mRNAs) granules per cell for a number of endogenous mRNAs. The grey box and line represent the interquartile range and the median, respectively. Each dot represents a single cell. n>300. (C) Beeswarm plot showing the proportion of mRNA that resides within multi-mRNA granules (>2.5mRNAs) per cell. Grey box and line represent the interquartile range and the median, respectively. n>300.

### Glycolytic mRNAs colocalize to the same RNA granules

Our previous assessment of the *ENO2* and *PDC1* mRNAs suggested that these two mRNAs colocalize to the same set of granules (Lui et al., 2014)(Figure 3A), but are distinct from the translation factor mRNA granules we have recently described as factories for the production and inheritance of the translation machinery (Pizzinga et al., 2019). On this basis, we speculated that the colocalization of glycolytic mRNAs might generally allow a concerted production and/or regulation of the pathway of the glycolytic enzymes. More recent work highlights the potential for co-translational assembly of components of the glycolytic pathway (Shiber et al., 2018), which again hints that actively translating glycolytic mRNAs might colocalize. In both the live cell and fixed cell mRNA localization experiments presented here, both the pattern and number of mRNA granules in a cell is remarkably similar across the different glycolytic mRNAs (cf. Figure 1C and 2A; cf Figure 1D and 2B). This similarity is consistent with a model where many of the glycolytic mRNAs colocalize to the same site.

**Figure 3.**
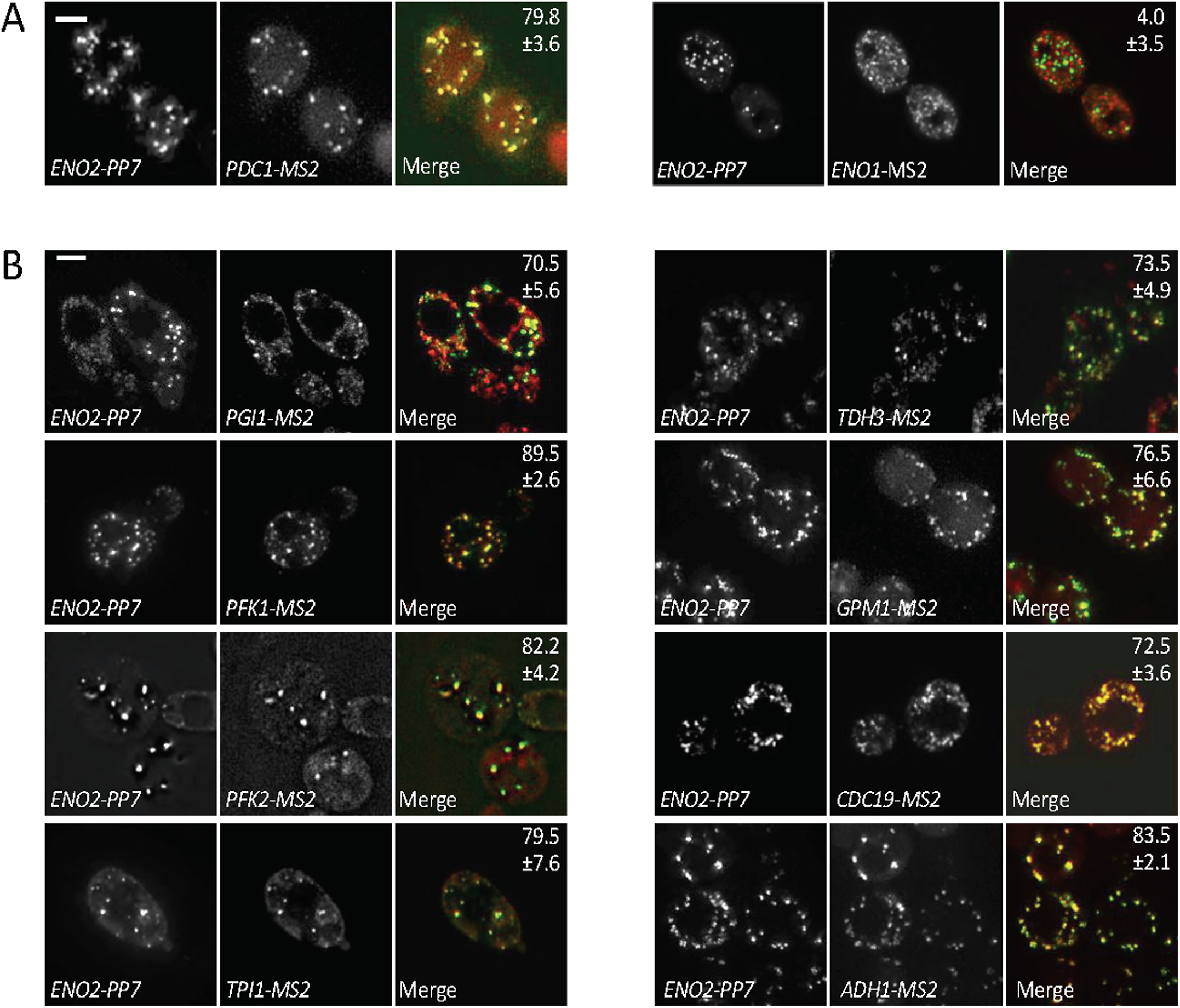
Glycolytic mRNAs colocalize to granules in actively growing cells. (A) and (B). *z*-stacked images show the localization of various *MS2*-tagged mRNAs (via co-expression of the MS2-CP-mCh fusion) relative to the *ENO2-PP7* mRNA (visualized using co-expression of the PP7-CP-GFP fusion). The percentage of observable tagged mRNA colocalizing with the *PP7*-tagged mRNA is indicated ± SD. Scale bars: 2 μm.

In order to directly assess glycolytic mRNA colocalization, we made use of a PP7/MS2 system, which allows the simultaneous visualization of two mRNAs in the same live cell (Hocine et al., 2013; Lui et al., 2014; Pizzinga et al., 2019). A series of yeast strains were generated carrying PP7-tagged *ENO2* mRNA and another MS2-tagged glycolytic mRNA. By co-expressing the MS2 and PP7 coat proteins fused to mCherry and GFP respectively, the localization of each MS2-tagged mRNA was compared directly to that of *ENO2* mRNA in the same living cell. As previously shown (Lui et al., 2014), we observed a strong colocalization of *PDC1* with *ENO2* using this system (Figure 3A). Equally, for many of the glycolytic enzymes tested, a high degree of colocalization with the *ENO2* mRNA pattern was observed (Figure 3B). Interestingly however, despite the high sequence homology between *ENO2* and *ENO1* mRNAs, these mRNAs localized to discrete foci (Figure 3A).

In order to corroborate the colocalization observed using the PP7/ MS2 systems, we assessed pairwise colocalization of endogenous unmodified glycolytic mRNAs using smFISH (Figure 4A). Since, in contrast to the MS2 system where only multi-mRNA granules can be followed, the smFISH technique detects all of the probed mRNA present in a cell, the signal is very congested. This congestion is particularly apparent for glycolytic mRNAs, which are typically estimated to be present at >100 copies per cell (Lahtvee et al., 2017). So, in order to objectively measure pairwise colocalization, a computational strategy was developed. In short, the distance between the centroid of a granule for one mRNA species was measured relative to the centroid of the nearest neighbouring granule for the other mRNA species. A spot was deemed to co-localize if its centroid was within the sphere of a spot in the opposite channel (Figure 4B). Using this method, a significant proportion (~50-60%) of the glycolytic mRNA granules were deemed to overlap with one another (Figure 4C). Since, each mRNA is present in ~20 multi-mRNA foci per cell (Figure 2B) with a similar number of single mRNA foci, we were concerned that high levels of colocalization could simply stem from the proportion of cytosolic space occupied by the mRNA foci. Therefore, to control for this, we established a Monte Carlo simulation model (Fletcher et al., 2010), where the position of real foci were randomized within the cell and cross-compared with randomized simulated foci for a second mRNA. From this analysis it is clear that relative to the simulated model, the various tested glycolytic mRNAs display significant colocalization (Figure 4C). While, these smFISH results mirrors the co-localization observed using the MS2 and PP7 stem loop systems (Figure 3), the scale of co-localisation appears lower. One possible reason for the lower colocalization reflects the extra sensitivity of the smFISH technique in detecting single mRNA foci, which may not co-localise to the same extent as multi -mRNA granules. To explore this, we considered only the most intense foci in the smFISH data, which likely represent multi-mRNA containing foci. This analysis revealed a positive shift in the level of co-localization by ~10-20% (Figure 4C). Importantly, simulated controls using these brighter, larger spots did not display this shift in co-localization (Figure 4C).

**Figure 4.**
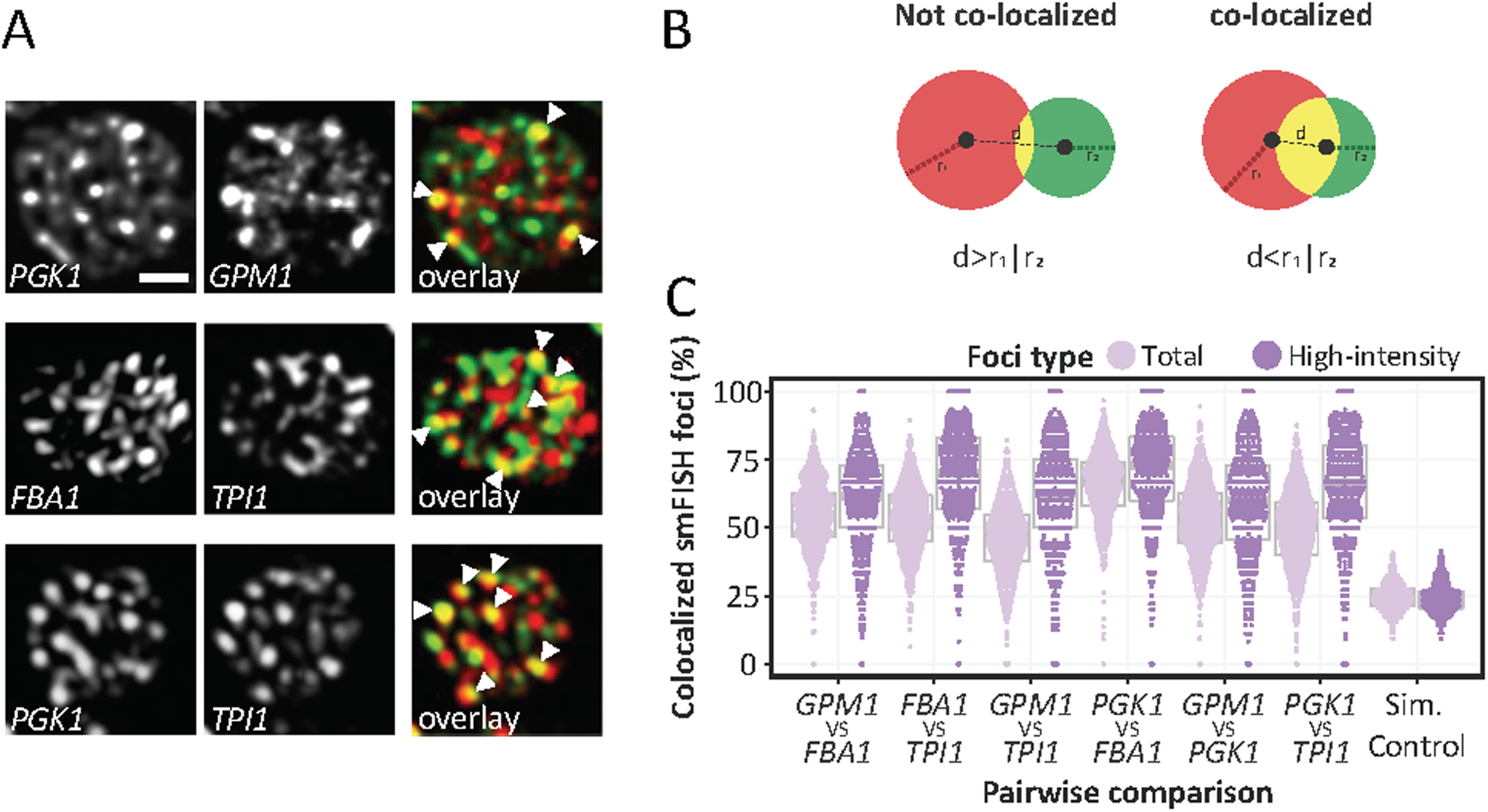
smFISH confirms that glycolytic mRNAs colocalize in granules. (A) Example *z*-stacked images from smFISH colocalization studies using the designated probes. Scale bar: 1μm. (B) Diagram detailing the colocalization method used to generate data in panel C. The centroid of both spots must be within the radius of either channel in order for spots to be deemed a colocalized (d<r_1_|r_2_). Spots that are touching are not always co-localized, if the distance between centroids is greater than the radius of both channels (d>r_1_|r_2_) (C) Beeswarm plot showing the proportion of colocalized smFISH foci considering total foci or only high-intensity foci, as indicated. Colocalization was assessed in a pairwise manner using smFISH foci identified via Fish-quant (see methods). Simulated colocalization was assessed by sub-sampling foci properties across a number of pairwise comparisons (see methods). Grey box and line represent the interquartile range and the median, respectively. Each data point represents a single cell, n>300.

Overall, the smFISH results combined with the live cell studies reveal that there is a high degree colocalization for the majority of glycolytic mRNAs tested to multi-mRNA granules in optimally growing yeast. Not every localized glycolytic mRNA co-localizes, for instance the *ENO1* mRNA does not, but the vast majority are. These colocalized mRNAs encode enzymes that catalyse the most of the reactions that are required for glucose fermentation to ethanol; therefore, we have termed the granules ‘<u>Co</u>re <u>Fe</u>rmentation’ mRNA granules or CoFe granules.

### mRNA translation both occurs in and is required for localization to CoFe granules

Previously, we have shown that in contrast to most mRNA containing granules, which carry translationally repressed mRNAs, the granules housing the glycolytic mRNAs *PDC1* and *ENO2* are sites where these mRNAs are translated (Lui et al., 2014). A variety of experiments supported this hypothesis, including data from FRAP assays where newly synthesized unbleached protein accumulated in the granules (Lui et al., 2014). In addition, treatment of cells with cycloheximide, which freezes ribosomes on mRNA, caused an increase in the number of mRNA granules. Furthermore, the quantification of ribosome-associated mRNA relative to granule-associated mRNA showed that even though 95% of these mRNAs are translated, at least 70% are present in granules. Finally, under polysome run-off conditions a rapid coalescence of the mRNA granules to form PBs was observed, suggesting that prior to the stress the mRNAs were in a distinct granule, being actively translated.

To extend this analysis further, a technique called TRICK (translating RNA imaging by coat protein knockoff) was used, which allows visualization of translation in live cells (Halstead et al., 2015; Pizzinga et al., 2019). This technique relies upon observations that a PP7 coat protein fusion bound to PP7 stem loops upstream of the STOP codon is displaced under active translation conditions, whereas the MS2 coat protein fusion tethered downstream of the STOP codon remains associated (Halstead et al., 2015) (Figure 5A). For *PDC1* mRNA under active growth conditions, most granules observed only carry the MS2-CP-mCherry (MS2-CP-mCh) fusion protein (Figure 5B and C). In contrast, after a 10 minute glucose depletion to elicit a robust and global inhibition of protein synthesis (Ashe et al., 2000), both MS2-CP-mCh and PP7-CP-GFP colocalize to granules (Figure 5B and C). This result supports our previous work showing that under active growth conditions the glycolytic mRNAs such as *PDC1* and *ENO1* are translated in granules, and combined with the colocalization studies, suggest that the CoFe granules serve as factories for glycolytic enzyme production.

**Figure 5.**
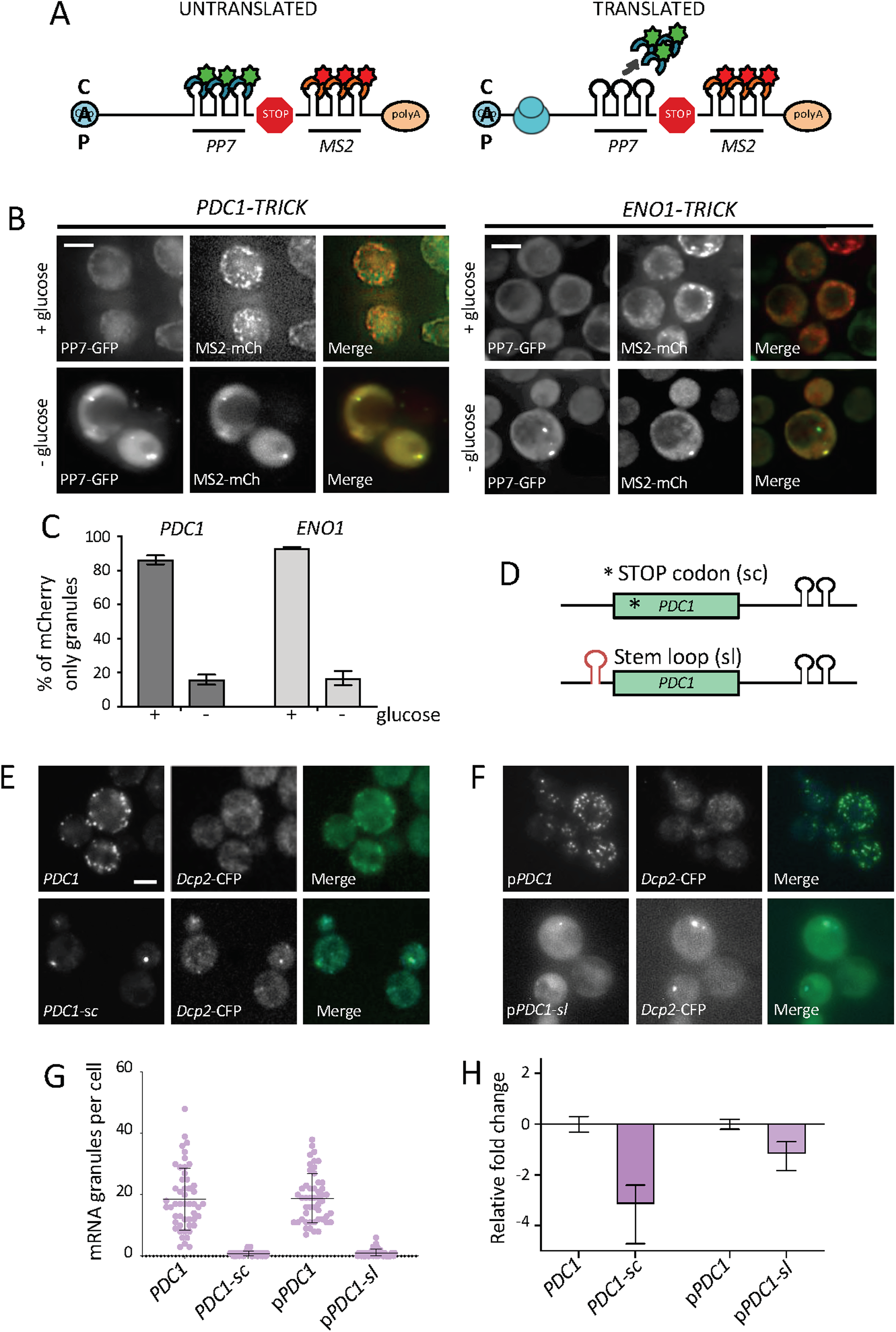
mRNAs translation in CoFe granules is required for localization. (A) Schematic of TRICK reporter system. Ribosomes on translated RNAs ‘knock off’ the PP7-CP-GFP fusion whilst on untranslated RNAs the coat protein remains bound. (B) *z* -stacked images of TRICK tagged mRNAs co-expressing the MS2-CP-mCh fusion and the PP7-CP-GFP fusion, in + and – glucose. Scale bars: 3 µm (C) Quantification of MS2-CP-mCh only granules as a percentage of total granules observed in TRICK tagged mRNAs in + and – glucose conditions. Error bars are ± SD. (D) Schematic of *PDC1* premature stop codon (sc) and stem loop (sl) insertion. (E) *z*-stacked images of cells expressing Dcp2p-CFP and *PDC1*-*MS2* tagged mRNA. *PDC1*-*MS2* (sc) has a premature stop codon in the ORF. (F) *z*-stacked images of strains expressing Dcp2p-CFP with p*PDC1*-*MS2* or p*PDC1*-*MS2* (sl). p*PDC1*-*MS2* (sl) has a stem loop upstream of the ORF. (G) Scatterplot of mRNA granules per cell in *PDC1*-*MS2* tagged mRNA with or without a premature stop codon and in strains bearing p*PDC1*-*MS2* with or without the stem loop. Error bars are ± SD. Scale bars: 2µm (H) Relative fold change of (i) *PDC1 MS2*-tagged mRNA relative to untagged *PDC1* mRNA, (ii) *PDC1*-*MS2* mRNA in strains harbouring a premature stop codon (sc) relative to a strain without, (iii) *PDC1*-*MS2* mRNA on a plasmid (p*PDC1*-*MS2* mRNA) relative to genomic *PDC1*-*MS2* mRNA and finally (iv) p*PDC1*-MS2 mRNA in strain with a stem loop upstream of the ORF relative to a p*PDC1*-*MS2* mRNA without a stem loop. Error bars are ± SD.

Another key question is whether translation of a molecule of mRNA is a requirement for entry into the granule. In order to address this question, we selected the *PDC1* mRNA and sought to limit its translation, then assess the impact on localization. More specifically, we adopted two different strategies toward reducing *PDC1* mRNA translation. In the first approach a STOP codon was inserted immediately downstream of the translation START codon (*PDC1-sc*) (Figure 5D). We reasoned that this would severely reduce the number of ribosomes associated with this mRNA and significantly increase the pool of non-translated *PDC1* mRNA. As a second strategy, a stem loop was inserted into the *PDC1* mRNA 5’UTR, upstream of the START codon (*PDC1-sl*) (Figure 5D). Introduction of this well-characterized stem loop (ΔG value of −41 kcal/mol) has previously been shown to reduce translation of specific mRNAs by limiting scanning of the 43S preinitiation complex through to the AUG Start codon (Palam et al., 2011; Pizzinga et al., 2019; Vattem and Wek, 2004). In strains carrying these altered *PDC1* mRNAs, mRNA localization was followed relative to the non -modified mRNA using the MS2 system.

Introduction of either the STOP codon or the stem loop structure into the *PDC1* mRNA dramatically reduced the number of *PDC1-MS2* mRNA granules: decreasing from ~20 granules per cell to less than 5 (Figure 5E, F and G). Coincident with this effect on the number of mRNA granules in the cell, both strategies used to limit *PDC1* mRNA translation also resulted in reduced mRNA levels (Figure 5H). Insertion of the STOP codon caused an ~8-fold reduction in *PDC1-MS2* mRNA, whereas stem loop insertion reduced mRNA levels ~2-fold. The reduction of mRNA caused by the introduction of the STOP codon is consistent with premature STOP codon mRNAs leading to nonsense mediated mRNA decay (Hagan et al., 1995). The impact of the stem loop on *PDC1* mRNA levels is not as pronounced as the STOP codon insertion, and it is a little surprising that this insertion leads to mRNA destabilisation, as this same stem loop has been inserted into a number of mRNAs without impacting upon overall mRNA levels (Palam et al., 2011; Pizzinga et al., 2019; Vattem and Wek, 2004). However, this result does highlight the intimate connection between the translation of an mRNA and its stability, and adds to many observations showing that a reduction in translation can lead to mRNA destabilization (Roy and Jacobson, 2013).

A surprising observation was made when the localization of the PB marker Dcp2p was assessed in cells bearing either the *PDC1-sc* or *PDC1-sl* mRNAs. In both strains, Dcp2p was constitutively present in PBs even in unstressed cells, and the *PDC1-sc* and *PDC1-sl* mRNAs colocalized with these bodies (Figure 5B). This result is especially intriguing as generally PBs are barely visible unless cells are stressed in some way (Lui et al., 2014). Yet in these unstressed cells, just a single point mutation to introduce a STOP codon into one mRNA species is sufficient to induce PB formation. This result is also interesting with regard to the controversy surrounding MS2-tagging. The specific introduction of a mutation that inhibits translation to destabilize an mRNA changes the pattern dramatically and causes PB formation. Therefore, if the RNA granules observed for non-mutated glycolytic mRNAs during exponential growth (Figure 1C) were due to the accumulation of RNA fragments carrying the MS2 stem loops, we would expect a similar colocalization with PB markers. However, we do not observe such colocalization with PB markers in unstressed cells (Lui et al., 2014; Pizzinga et al., 2019).

Overall, these results highlight that in keeping with many observations over the years it is difficult to alter the translation of an mRNA without affecting its stability (Mugridge et al., 2018; Roy and Jacobson, 2013). However, the results do suggest that as well as translation occurring in the glycolytic mRNA granules, translation is important for mRNAs to enter these granules, since when translation is reduced alternative mRNA fates such as degradation and relocalization to PBs become apparent.

### Glycolytic mRNA localization varies according to the level of fermentation

In order to understand the potential physiological role the CoFe granules play, a series of experiments were undertaken where yeast were grown on a range of carbon sources selected based upon the pathways required for carbon source metabolism. For example, while yeast cells ferment glucose to ethanol even under aerobic conditions, for other carbon sources the degree of fermentation varies. Yeast cells grown on ethanol as the sole carbon source derive their energy from respiration and only require the glycolytic enzymes for gluconeogenesis. Raffinose is catabolised initially via the action of the secreted enzymes invertase (Suc2p) and α-galactosidase (Mel1p). These enzymes yield the monosaccharides glucose, fructose and galactose, which are readily available for fermentation via the glycolytic enzymes (Barnett, 1976). Equally, yeast that are pre-adapted to galactose, express enzymes of the Leloir pathway allowing fermentation of this sugar via entry into the glycolytic pathway (Timson, 2007).

Microscopic analysis revealed that yeast cells grown on glucose, raffinose or galactose harboured approximately 10-20 granules of either *PDC1* mRNA or *ENO1* mRNA per cell, whereas cells grown on ethanol harboured significantly fewer granules (Figure 6A and B). In terms of the levels of the *PDC1* and *ENO1* mRNAs, these also vary with carbon source, consistent with glucose representing the preferred yeast carbon source. For both mRNAs, glucose grown cells harbour significantly more glycolytic mRNA than cells grown on most other carbon sources (Figure 6C). Therefore, these results show that both the level of glycolytic mRNAs and prevalence of CoFe granules vary depending on the carbon sources. In particular, the presence of CoFe granules appears to correlate with a requirement for glycolytic flux to utilise the provided carbon source. In contrast, the overall level of the mRNA is more tied to the quality of the carbon source rather than the pathway required for its utilization.

**Figure 6.**
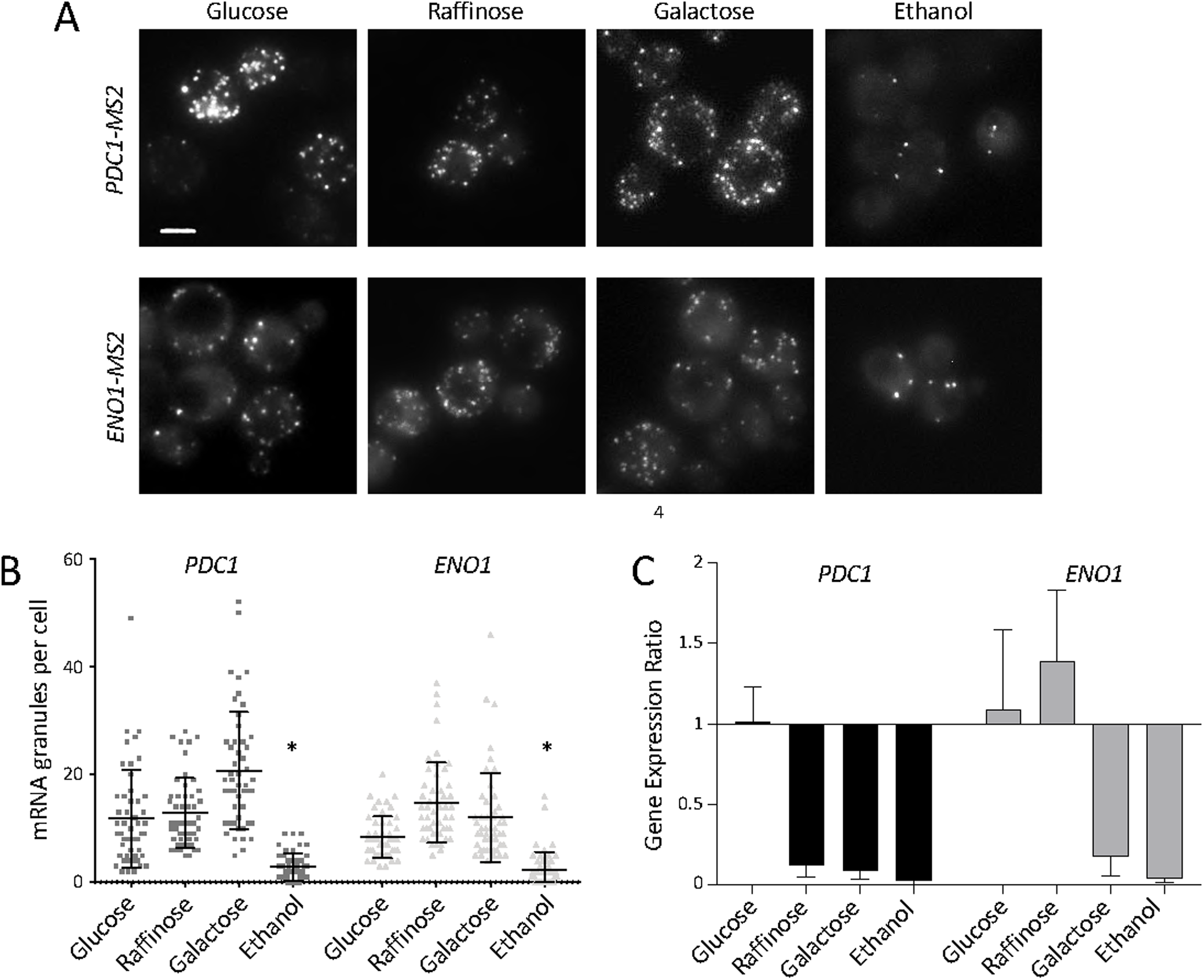
CoFe granule number varies with quality of carbon source. (A) *z*-stacked images of *PDC1-MS2* or *ENO1-MS2* mRNA in strains co-expressing the MS2-CP-GFP fusion, grown in SC media with either 2% glucose, 2% raffinose, 2% galactose or 3% ethanol. Scale bar: 2μm. (B) Quantification of mRNA granules per cell for *PDC1* and *ENO1* mRNA in strains grown in the different carbon sources. n=50. Error bars are ± SD (C) Graph representing the gene expression ratio of *PDC1* or *ENO1* in strains (yMK1586 and yMK2468) grown in the different carbon sources. The gene expression ratio was calculated according to the Pfaffl method. This approach considers the PCR efficiency for the different genes and is expressed as the change in target gene levels (*PDC1* and *ENO1*) between glucose conditions and the different carbon sources over the change in reference gene levels (*ACT1*) between glucose conditions and the different carbon sources (Pfaffl, 2001). Error bars are ± SD.

Overall, the data are consistent with a view that the localization of glycolytic mRNAs to CoFe granules represents a strategy allowing high-level co-ordinated production of glycolytic enzymes in translation factories.

### Glycolytic mRNA granules are also evident in human cells

In order to assess whether a similar organization of glycolytic mRNAs might exist in higher eukaryotic cells, smFISH analysis was conducted for four different glycolytic mRNAs in HeLa cells: two enolase mRNAs (*ENO1* and *ENO2*), lactate dehydrogenase mRNA (*LDHA)* and a phosphofructokinase mRNA (*PFKM)*. For all four of the selected mRNAs, variation in mRNA signal was observed both in terms of particle size and intensity (Figure 7A, data not shown). In particular large intense mRNA foci were observed suggesting that granules harbouring multiple mRNAs can also be a feature of higher eukaryotic cells. Similar observations were made in other cell lines such as HFF-1 cells and SH-SY5Y cells (data not shown). This opens up the possibility that glycolytic mRNAs might be co-ordinately localized in higher cells. Therefore, a multichannel smFISH approach was taken. Here, evidence for a specific colocalization of the *ENO2* and *PFKM* mRNAs was obtained (Figure 7B and C). While the degree of colocalization is not as comprehensive as observed in yeast, these data do show that in actively growing human tissue culture cells, glycolytic mRNAs can be localized to granules, and that these granules can contain more than one type of glycolytic mRNA.

**Figure 7.**
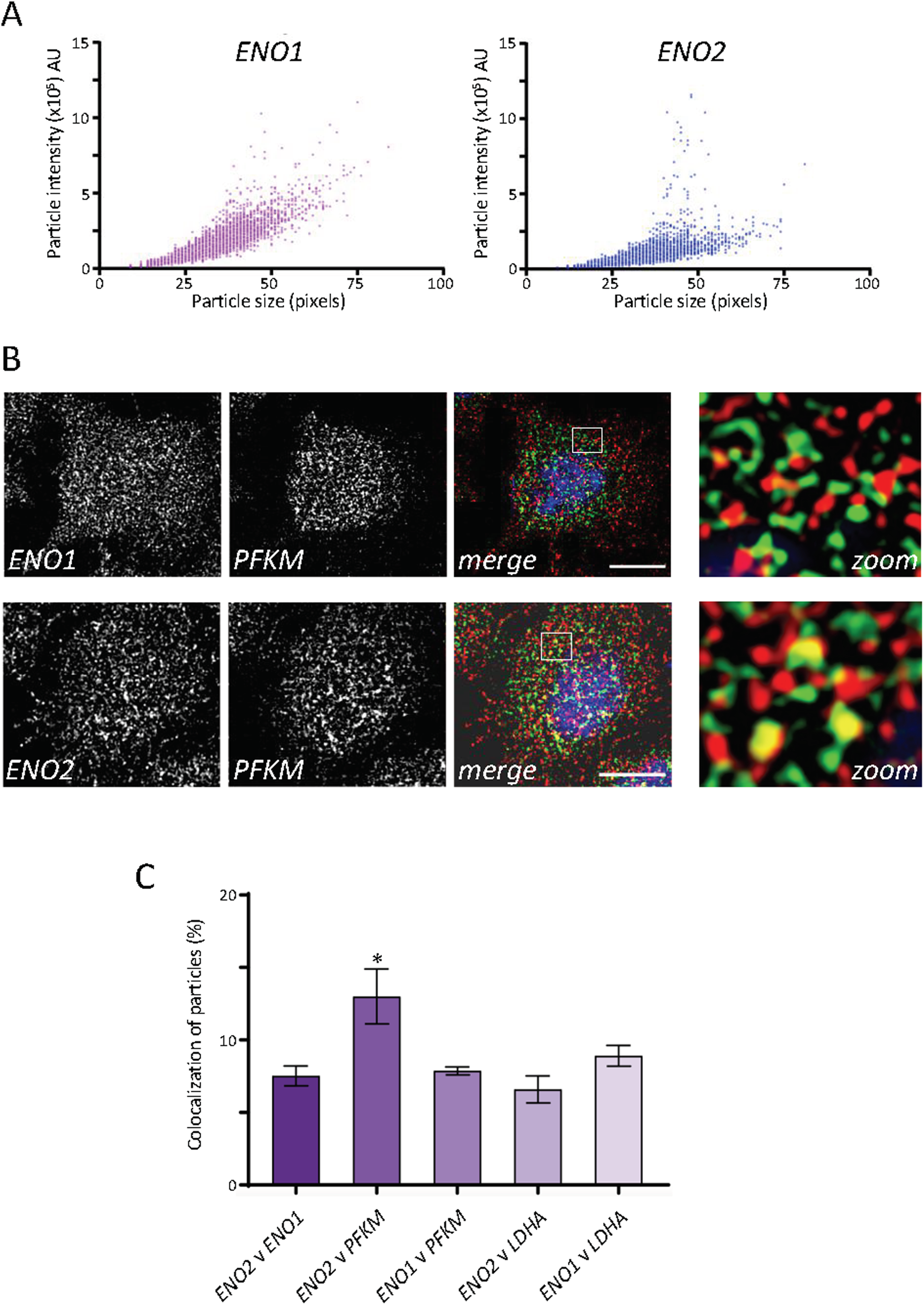
Human glycolytic mRNAs are present in granules and can colocalize. (A) Scatter plot of particles detected in *z*-stacked smFISH images of Hela cells using probes to the mRNAs indicated. Images from 3 biological replicates were analyzed using the Image J ComDet plugin. Particle size is measured in pixels where each pixel = 45×45 nm. (B) Single *z* -slice smFISH images of HeLa cells using probes to the mRNAs indicated. Scale bar: 10μm. Insets magnified x2. (C) Histogram showing the percentage of colocalized mRNA particles calculated using ComDet analysis of *z*-slices from 3 biological replicates. The significance across the various combinations was calculated using one-way ANOVA. *ENO2* v *PFKM* (P value < 0.05 shown by asterisk) is significantly different to *ENO2* v *ENO1*, *ENO1* v *PFKM* or *ENO2* v *LDHA*.

## Discussion

mRNA localization serves critical functions in the expression of proteins at specific loci within cells and in the response to stress in terms of PB and SG formation (Pizzinga and Ashe, 2014). In this study, we suggest that mRNA localization to granules can co-ordinate whole pathways of metabolism. We use a combination of live cell experiments and smFISH to show that glycolytic mRNAs localize to granules in yeast and human cells. In stark contrast to mRNAs localizing to PBs, SGs or transport granules, in yeast the glycolytic mRNAs are translated in CoFe granules, and their translation is a requirement for localization.

Recent evidence suggests that liquid-liquid phase separation (LLPS) within cells produces membraneless compartments or biological condensates where enzymatic reactions and processes can occur. For instance, in the nucleolus, rRNA is produced via numerous highly complex reactions (Brangwynne et al., 2011), while in the centrosome microtubule nucleation occurs (Zwicker et al., 2014). The CoFe granules described here conform to many of the properties of phase-separated condensates: they are dynamic, can be observed to fuse and are disrupted by low concentrations of 1,6-hexanediol (Lui et al., 2014). Therefore, our data suggest that translated glycolytic mRNAs are present in such biological condensates where molecular processes are not only maintained but might actually be enhanced (Kojima and Takayama, 2018).

Enhanced translation of mRNA is therefore one possible explanation as to why the glycolytic mRNAs would be localized within granules. Previous observations from our lab have shown that up to 95% of the glycolytic mRNAs are translated (Lui et al., 2014). In addition, the glycolytic mRNAs are amongst the most abundant in the cell and so may require rather specific mechanisms to maintain their high rates of translation. Equally LLPS has previously been associated with altered efficiency of a host of biological processes and enzymes (Zhou et al., 2008), so translation may prove to represent another example of such a process, especially where the co-ordinated generation of high volumes of glycolytic enzyme may be important.

Another possible rationale for localized mRNA translation is to aid the formation of multi-protein complexes. Many of the glycolytic enzymes are present in multimeric complexes. For example, almost all of the glycolytic enzymes function as multimers: in yeast the phosphofructokinase enzyme is present as an octamer (Schwock et al., 2004), phosphoglycerate mutase and pyruvate kinase are tetramers (Jurica et al., 1998; Rigden et al., 1998), and enolase is dimeric (Sims et al., 2006). Co-translation of individual mRNAs at the same site within cells could therefore aid the formation and productive folding pathways for these complexes. A range of precedents exist for the co-translational production of complexes across various biological systems (Halbach et al., 2009; Kamenova et al., 2019; Shiber et al., 2018; Wells et al., 2015), while a systematic analysis in *Schizosaccharomyces pombe* suggests that co-translational production of protein complexes is widespread, with a substantial fraction of proteins co-purifying with mRNAs that encode interacting proteins (Duncan and Mata, 2011). Notably, in recent work characterizing the propensity for co-translational folding in several different protein complexes in *Saccharomyces cerevisiae*, the *PFK1* and *PFK2* phosphofructokinase mRNAs were identified as key examples where the translated products are co-translationally assembled or folded (Shiber et al., 2018).

It has also been shown that, as well as forming multimeric single enzyme complexes, various different glycolytic enzymes can be compartmentalized into much larger complexes (Masters, 1991). A variety of observations suggest that the physical compartmentalization of glycolysis is advantageous. For instance, in protozoan organisms such as *Trypanosoma* and *Leishmania*, a specific membrane-bound organelle called the glycosome has evolved, which is thought to provide these pathogens scope for regulating metabolic activity (Haanstra et al., 2016). Furthermore, in human cells, such as skeletal muscle cells, neurons and erythrocytes; glycolytic enzymes can be organized as complexes co-ordinated either on membranes or the cytoskeleton (Knull and Walsh, 1992; Puchulu-Campanella et al., 2013). Moreover, a glycolytic metabolon has also been described in yeast (Masters, 1991) and it is thought to be stabilized by various weak interactions with actin (Araiza-Olivera et al., 2013). These multi-enzyme complexes are likely to promote both the channelling of metabolites from one enzyme to the next, as well as the reduction of potentially toxic intermediates. More recent work in yeast has shown that while glycolytic enzymes are broadly cytosolic under non-stress conditions, they can coalesce into ‘G-bodies’ in response to hypoxic stress (Jin et al., 2017). Overall therefore, the co-production of the glycolytic enzymes at the same site by virtue of co-ordinated mRNA localization could promote the co-translational formation of some of these higher order complexes of enzymes. Although it should be noted that our own data suggest that under active growth conditions fluorescent-protein tagged forms of the glycolytic enzymes are generally found throughout the cytosol (Lui et al., 2014).

Another point worth reflecting upon when considering the role of the CoFe granules is that several glycolytic enzymes have extra-glycolytic or ‘moonlighting’ functions outside of their role in glycolysis. For example, many of the glycolytic enzymes have been identified as RNA binding proteins that appear to interact with their own mRNA (Castello et al., 2015; Matia-Gonzalez et al., 2015). In addition, yeast enolase is important for both the mitochondrial import of tRNA Lys CUU (Entelis et al., 2006) and for vacuole fusion (Decker and Wickner, 2006), and yeast fructose-1,6-bisphosphate aldolase is important for vacuolar H+-ATPase function (Lu et al., 2007). Many further moonlighting functions of glycolytic enzymes, including nuclear functions in transcription, DNA replication/ repair and histone modification have been described (Boukouris et al., 2016). One possible explanation for the presence of CoFe granules could be that they serve as a focus for the co-ordinated high-level production of the glycolytic machinery *en masse*, whereas individual translated glycolytic mRNAs outside of factories could provide the capacity for moonlighting protein activities.

One intriguing observation made during the course of our studies, is that while the yeast *ENO1* mRNA is observed to localise to granules, it does not appear to colocalize with the *ENO2* mRNA in CoFe granules. Several possible non-mutually exclusive explanations could account for this. Firstly, the expression levels of the enolase isoforms vary greatly depending upon the fermentation/ respiration status; Eno2p represents the predominant polypeptide in fermenting cells whereas the distribution is much more even in stationary phase or respiring cells (Entian et al., 1987). So the discrete localization of the two mRNAs could contribute to these expression differences. Secondly, since the enolase enzyme is dimeric, and both Eno2p and Eno1p homodimers, as well as the heterodimer have been described (Holland et al., 1982), the discrete localization of the *ENO1* and *ENO2* mRNAs could serve to regulate the relative proportion of these different complexes via cotranslational homodimer formation. The rationale for requiring homodimers seems unlikely to reside in their enzymatic function, as the various forms display very similar enzyme kinetics (Holland et al., 1982). Therefore, an alternative possibility relates to the enolase moonlighting functions, with Eno2p playing a more active role than Eno1p in the targeting of the nuclear-encoded tRNA^Lys^_CUU_ isoacceptor to mitochondria (Entelis et al., 2006).

Overall, glycolysis is perhaps the most fundamental of all biological pathways. The enzymes of the pathway are regulated at almost every level and paralogues have evolved distinct functions. The pathology of many disease conditions is intimately connected to the glycolytic pathway. For instance, aerobic glycolysis serves as a hallmark of many malignant cancers and the surrounding stroma, which can serve as a negative prognostic indicator due to increased resistance to therapy (Lee and Yoon, 2015; Ngo et al., 2015). The identification and characterization of factories for the production of glycolytic proteins can only serve to increase understanding of the functions, regulation and possibility for genetic adjustment of this key metabolic pathway.

## Acknowledgements

We thank L. Berchowitz, J. Gerst, J. Chao and R. Singer for reagents; P. March and S. Marsden for microscopy advice; and E. Linney for comments of the manuscript. FMP was supported by a CONICYT Becas Chile studentship (72140307). CB and MP were supported by Wellcome Trust (WT) PhD studentships (210002/Z/17/Z and 099732/Z/12/Z). JL, CG and GF were supported by a Biotechnology and Biological Sciences Research Council (BBSRC) project grant (BB/K005979/1). JC was supported by a BBSRC project grant (BB/P018270/1). The Bioimaging microscopes used in this study came from with grants from BBSRC, WT and the University of Manchester Strategic Fund.

## Author contributions

All authors generated reagents, performed experiments, evaluated results and generated figures. FMP, CB, JL, JC, MPA conceived the study and designed experiments, while MPA wrote the manuscript. All authors contributed to the discussion and evaluation of the manuscript.

## MATERIALS AND METHODS

### RESOURCE AVAILABILITY

#### Lead Contact

Further information and requests for resources and reagents should be directed to the Lead Contact, Mark P Ashe.

#### Materials Availability

Strains and plasmids generated as part of this study are available upon request.

#### Data and Code Availability

Code for bespoke yeast cell smFISH analysis is available at the github repository github.com/CPBS/MoralesPolanco

### EXPERIMENTAL MODEL AND SUBJECT DETAILS

#### Yeast growth conditions

Yeast experiments were performed in the *Saccharomyces cerevisiae* strain yMK467-a derivative of W303-1A (SCR_003093). Unless stated otherwise, experiments were performed after strains were grown in synthetic complete (SC) media with 2% glucose at 30°C to exponential phase. Strains used in this study are listed in Table I. For live-cell microscopy, cells were pelleted at 500xg for 3 minutes at 30°C, resuspended in pre-warmed (30°C) media lacking methionine and incubated for 30 min to induce expression of the pCP-GFP/mCh fusions prior to imaging. For growth on alternative carbon sources SC media was supplemented with 2% raffinose, 2% galactose or 3% ethanol. For stress conditions, cells were incubated in media lacking glucose for 10 minutes.

#### Human cell-line growth conditions

Human cell-line experiments were performed in HeLa cells (CVCL_0030), grown in Dulbecco’s modified Eagle’s medium supplemented with 10% fetal bovine serum at 37 °C.

### METHOD DETAILS

#### Yeast strain and plasmid construction

MS2 and PP7 tagged strains were generated as previously described (Haim-Vilmovsky and Gerst, 2011; Hocine et al., 2013; Tutucci et al., 2018), using plasmid reagents generously donated by Jeff Gerst and Robert Singer. Dual MS2 and PP7 tagged strains were generated by crossing the single tagged haploid strains. Subsequent diploid strains were selected, sporulated and the appropriate dual tagged haploid strains were verified by PCR. The strain harbouring a premature stop codon in the *PDC1* ORF was generated via recombination of a mutant PCR product generated from the *PDC1-MS2* tagged strain. More specifically, oligonucleotides with a specific mutation in the upstream primer were used to amplify a *PDC1::HIS5::MS2* cassette from genomic DNA prepared from an intermediate strain in the *PDC1-MS2* m-TAG procedure (Haim-Vilmovsky and Gerst, 2011). The mutation introduces a premature STOP codon in the *PDC1* ORF. The *PDC1::HIS5::MS2* cassette was then transformed and recombined into the *PDC1* locus of the yMK467 strain. Removal of the *HIS5* marker was carried out using a Cre recombinase strategy as previously described (Haim-Vilmovsky and Gerst, 2011). TRICK strains were generated using a similar approach to MS2 or PP7 tagging, but using a DNA template developed for TRICK in yeast (Pizzinga et al., 2019). For generation of the yEPlac195-*PDC1* (p*PDC1*-*MS2*) plasmid, *PDC1-MS2* was amplified from genomic DNA of yMK1586 and cloned into the pGEM -T Easy vector and subsequently cloned into yEPLac195 using *Sph*I and *Sac*I restriction enzymes. A stem loop sequence (Vattem and Wek, 2004) was inserted upstream of the start codon in *PDC1* using Gibson assembly (Gibson et al., 2009) to generate plasmid yEPlac195-PDC1-SL (p*PDC1*-MS2 (sl)).

#### Single molecule fluorescent *in situ* hybridisation (smFISH)

For yeast cultures, smFISH was performed as previously described (Pizzinga et al., 2019). In brief, exponential-phase yeast were fixed in 4% EM-grade formaldehyde (15714-S; Electron Microscopy Sciences) for 45 min, then spheroplasted and permeablized with 70% ethanol. Gene-specific 20nt antisense oligonucleotides were designed with a 59nt Flap sequence, to which fluorescently labelled oligonucleotides were annealed (Pizzinga et al., 2019; Tsanov et al., 2016). The conjugated fluorophores included Alexa Fluor 488, Alexa Fluor 546, ATTO 590 and Alexa Fluor 648. After careful titration of the probe, 20 pmol fluorescently labelled smFISH probe was found to generate optimal signal relative to background when added to the cells in hybridization buffer (10 mg E. coli tRNA, 2 mM Ribonucleoside Vanadyl Complex, 200 μg/ml BSA, 10% dextran sulfate, 10% formamide, and 2× SSC in nuclease-free water). This hybridization buffer – probe mix was then added to cells and incubated overnight at 37°C, with gentle agitation. Cells were then washed in 10% formamide and 2× SSC and adhered to 0.01% poly-L-lysine–coated coverslips before mounting in ProLong diamond antifade mounting solution with DAPI (Life Technologies Cat# P36970).

For human cell experiments, HeLa cells were seeded in Dulbecco’s modified Eagle’s medium supplemented with 10% fetal bovine serum onto 13 mm laminin coated coverslips in sterile 24-well plates, then were fixed in methanol for 10 min at −20oC. Fixed cells were washed in 10% Formamide, 2x SSC buffer in nuclease-free water for 30 min at room temperature, then hybridized probes (as above) were added at a concentration of 25 nM for Cy7-conjugated probes and 75 nM for Cy5-conjugated probes at 37 oC overnight. Cells were then washed and mounted (Tsanov et al., 2016). An extensive series of experiments was undertaken to optimise the probe concentration and fixation method such that the no signal was detected in the various channels when a particular probe was absent.

#### Live-cell fluorescent microscopy

All yeast live-cell epifluorescent microscopy was performed on a Nikon Eclipse E600 or a Delta Vision microscope (Applied Precision) equipped with a Coolsnap HQ camera (Photometrics), using a 100x/ 1.40 NA oil plan Apo objective. Fluorescent parameters for each fluorophore are as follows; GFP (excitation −490/20 nm, emission-535/50 nm); mCherry (excitation-572/35 nm, emission-632/60 nm); and CFP (excitation-436/10 nm, emission-465/30). For routine live-cell imaging, exponentially growing cells were viewed on poly-L-lysine coated glass slides and images were taken with a *z*-spacing of 0.2μm. Images were acquired using Softworx 1.1 (Applied Precision), or Metamporph (Molecular Devices) software and processed using Image J software package (National Institute of Health, NIH).

#### Fixed-cell fluorescent microscopy

Images of human cells were acquired on an Olympus IX83 inverted microscope using Lumencor LED excitation, a *100x* objective and the Penta filter set. The images were collected using a *Retiga R6 (Qimaging)* CCD camera with a *z*-optical spacing of 0.2 μm. Raw images were then deconvolved using the Huygens Professional software (Scientific Volume Imaging).

Yeast smFISH images were collected on a Leica TCS SP8 AOBS inverted gSTED microscope using a 100x/1.40 Plan APO objective and 1x confocal zoom, as described previously (Pizzinga et al., 2019). DAPI staining was detected using a photon multiplying tube with a blue diode 405nm laser (5%). Confocal images of smFISH signals were collected using hybrid detectors with the following detection mirror settings; Alexa Fluor 488 498-536nm; Alexa Fluor 546 556-637nm (5 to 35µs gating); ATTO 590 603-637nm; Alexa Fluor 647 657-765nm using the 488nm (60%), 550nm (60%), 593nm (60%) and 646nm (60%) excitation laser lines, respectively. Images were collected sequentially in 200nm *z-*sections. Acquired images were subsequently deconvolved and background subtracted using Huygens Professional (Scientific Volume Imaging).

#### Quantitative RT-PCR (qRT-PCR)

RNA preparations were carried out using Trizol as described by the manufacturer (Thermofisher scientific, Cat#15596026), followed by isopropanol precipitation then treatment with Turbo DNase (Thermofisher scientific, Cat#AM2238). qRT-PCR was performed in a two-step manner using a ProtoScript First Strand cDNA synthesis kit (New England Biolabs, Cat#E6300S) and iQ SYBR Green Supermix (Bio-Rad, Cat#1708880) according to manufacturer’s instructions. Reactions were performed using 100ng of cDNA. iTaq Universal SYBR Green One Step Kit (Bio-Rad, Cat#1725150) was used to carry out one-step qRT-PCR and reactions were performed using 300ng of RNA. A CFx Connect Real -Time system was used to run reactions. Samples were run in triplicate and normalized to *ACT1* mRNA, and the fold change was calculated using either the 2^-ΔΔCq^ or the Pfaffl method (Livak and Schmittgen, 2001; Pfaffl, 2001).

### QUANTIFICATION AND STATISTICAL ANALYSIS

#### Quantification of microscopy and statistics

For quantification of granule numbers per cell from live cell experiments, 50 cells were counted for each strain. For quantification of overlapping MS2 and PP7 signal in double-tagged strains or TRICK strains, 100−150 granules were considered for each strain over three biological repeats. GraphPad Prism 6 (GraphPad Software, Inc.) was used to produce the graphs and to calculate the standard deviation, indicated by error bars. Two-way ANOVA was performed using GraphPad Prism 6. * denotes a P value < 0.0001.

smFISH images were processed and analysed using FISH-quant (Mueller et al., 2013) and FindFoci (Herbert et al., 2014) to identify spot position and size and provide spot enhancement via dual Gaussian filtering. To account for differences in smFISH signal intensity between fluorophores and experiments, different intensity thresholds for each channel/image were determined manually. However, the same thresholds were applied to all cells in that image. Cell outlines were automatically generated using a modified version of the CellProfiler (Carpenter et al., 2006) pipeline provided with FISH-quant that utilizes background cytoplasmic DAPI staining rather than brightfield images to determine cytoplasmic cell boundaries. Spot colocalization and other foci characteristics were assessed and quantified using custom scripts in python to scrape data from FISHquant into long format and R for more detailed analysis. Colocalization analysis was performed by pairing spots between channels based on spot centroid distance in 3D space (Eliscovich et al., 2017). Spots were deemed to colocalize if the 3D distance between centroids was less than the radius of either of the two spots, i.e. if the centroid of one spot existed within the radius of another spot. mRNA quantitation was performed using Gaussian fitting, as described previously (Pizzinga et al., 2019). To account for stochasticity in initial fitting parameters, this fitting was repeated 1,000 times and averaged. Simulated controls, based on the Monte Carlo simulation method (Fletcher et al., 2010), were performed by randomly sub-sampling spot characteristics, such as size in *x*, *y* and *z* planes, and arbitrarily positioning these within a simulated volume, typical of a yeast cell, as measured using the custom CellProfiler pipeline. The colocalization of these randomly positioned foci was subsequently processed using the same script outlined above, and iterated 1,000 times per pairwise comparison.

### KEY RESOURCES TABLE

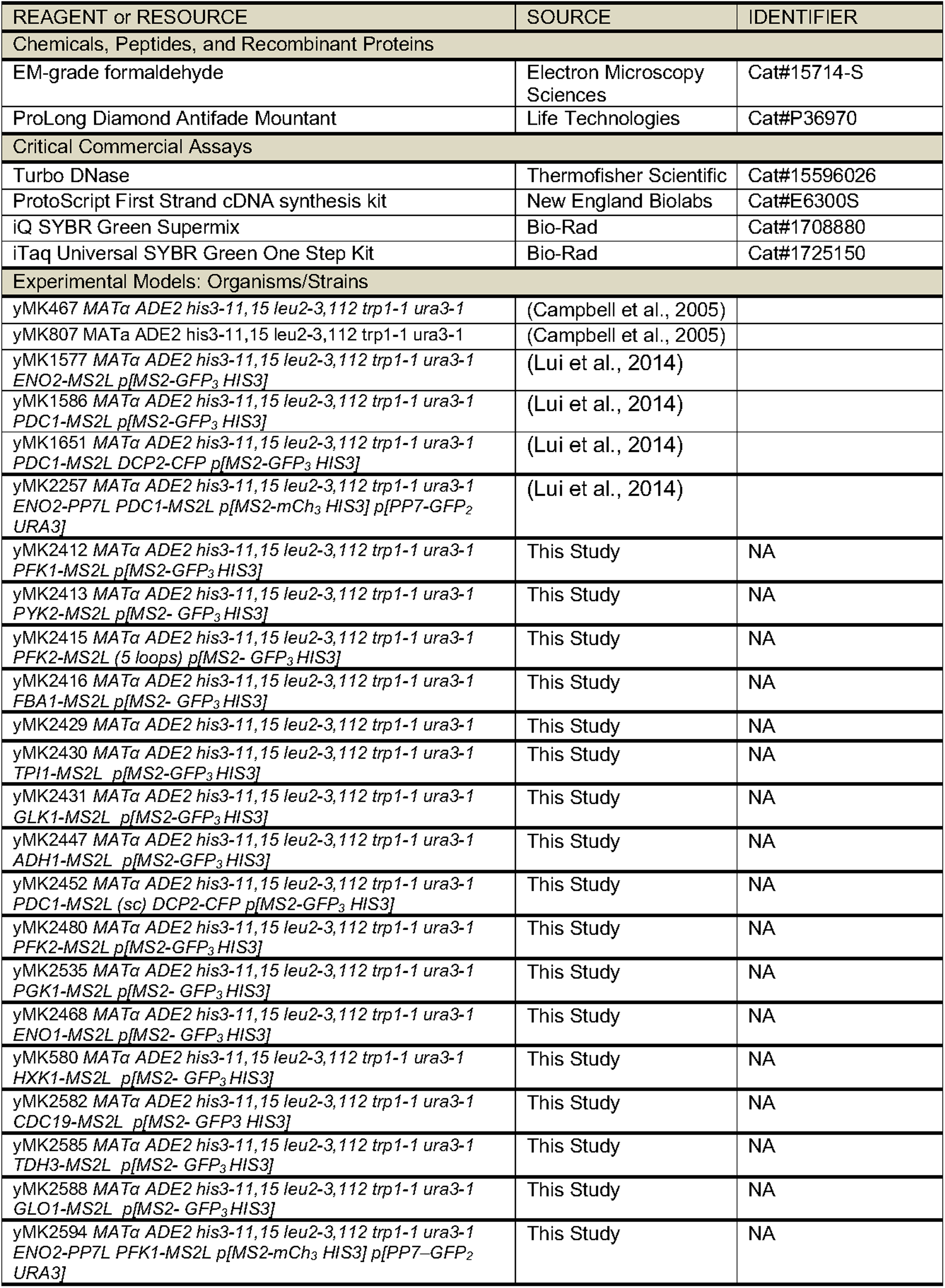
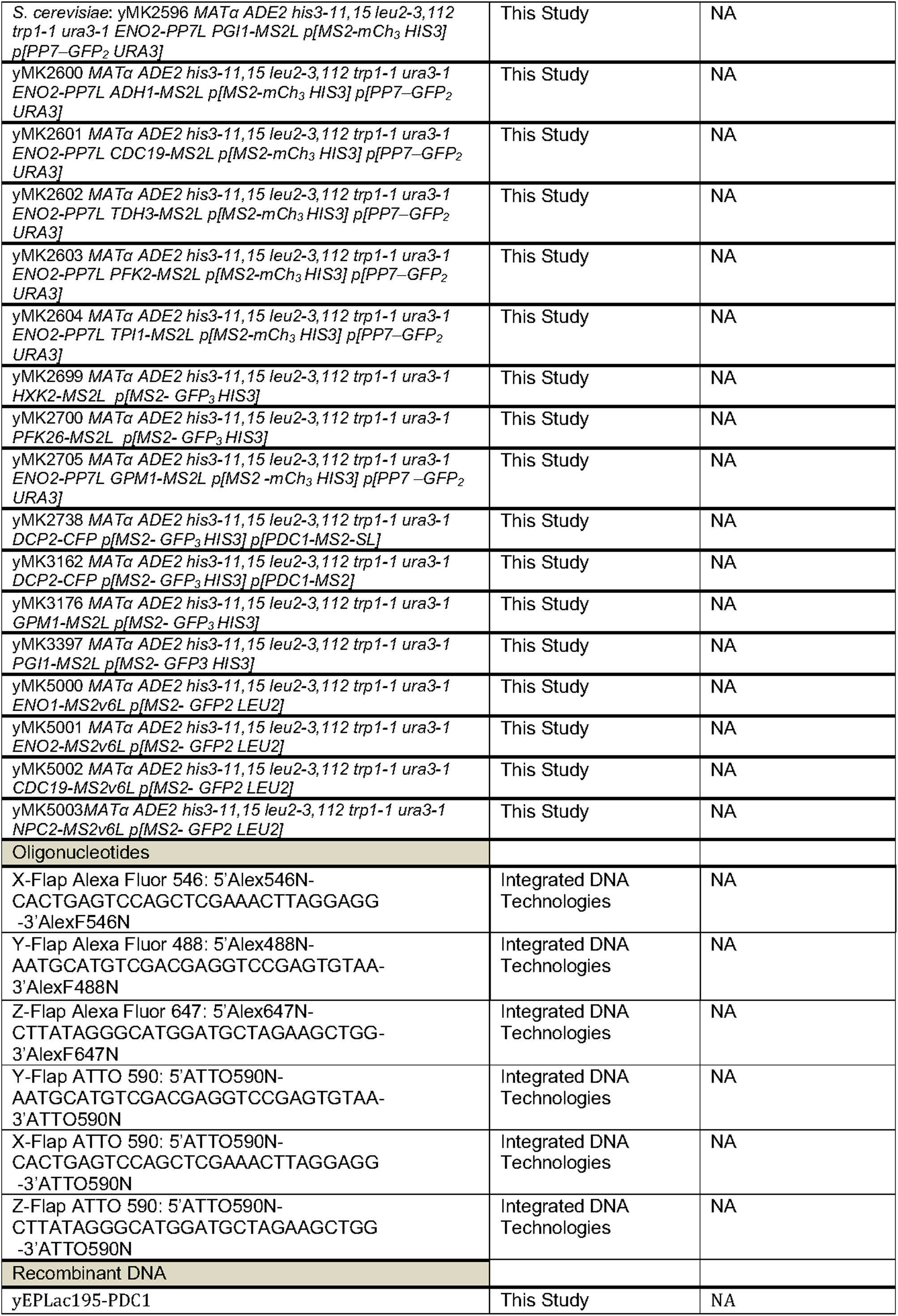
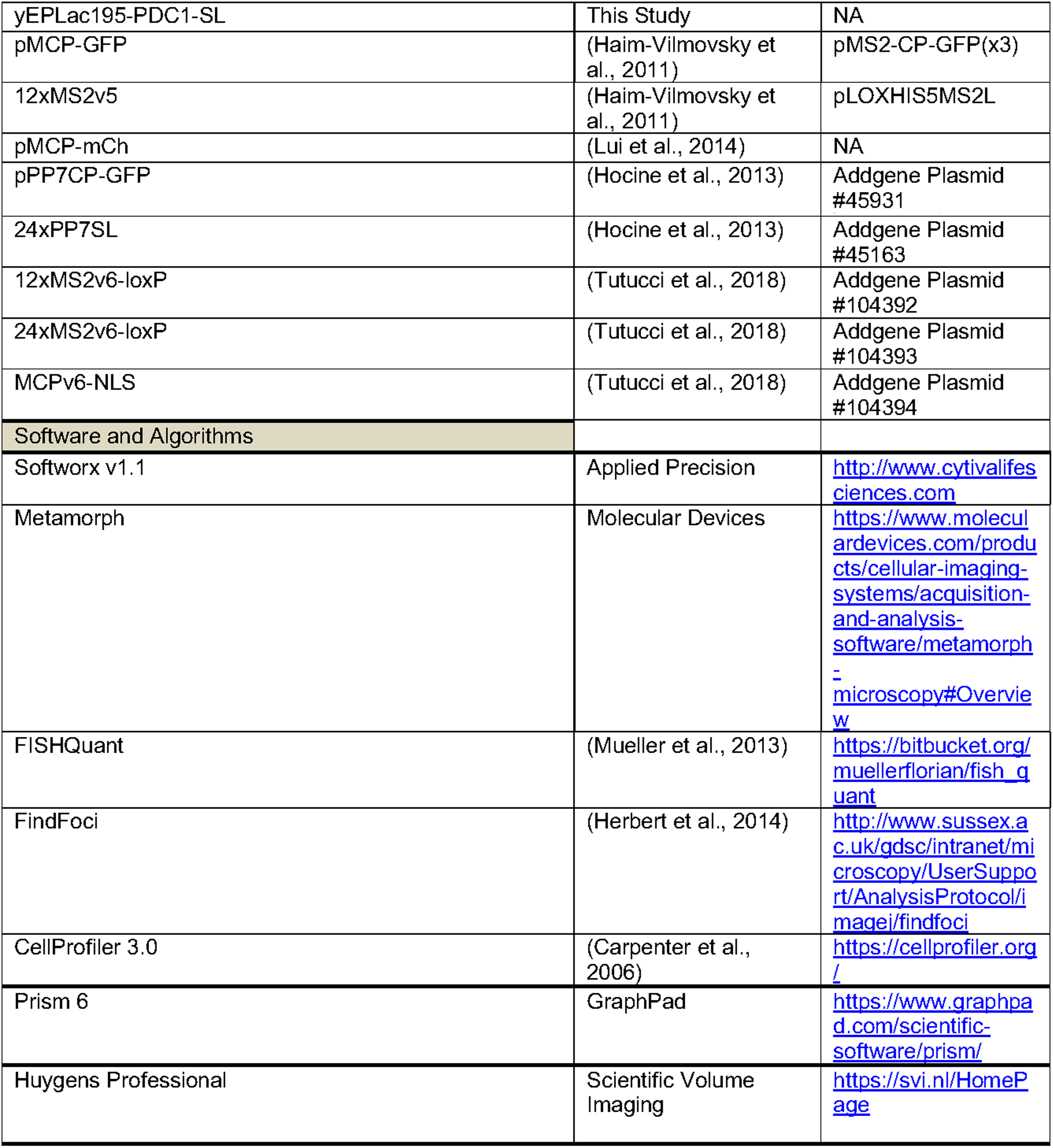

---

**Figure S1.**
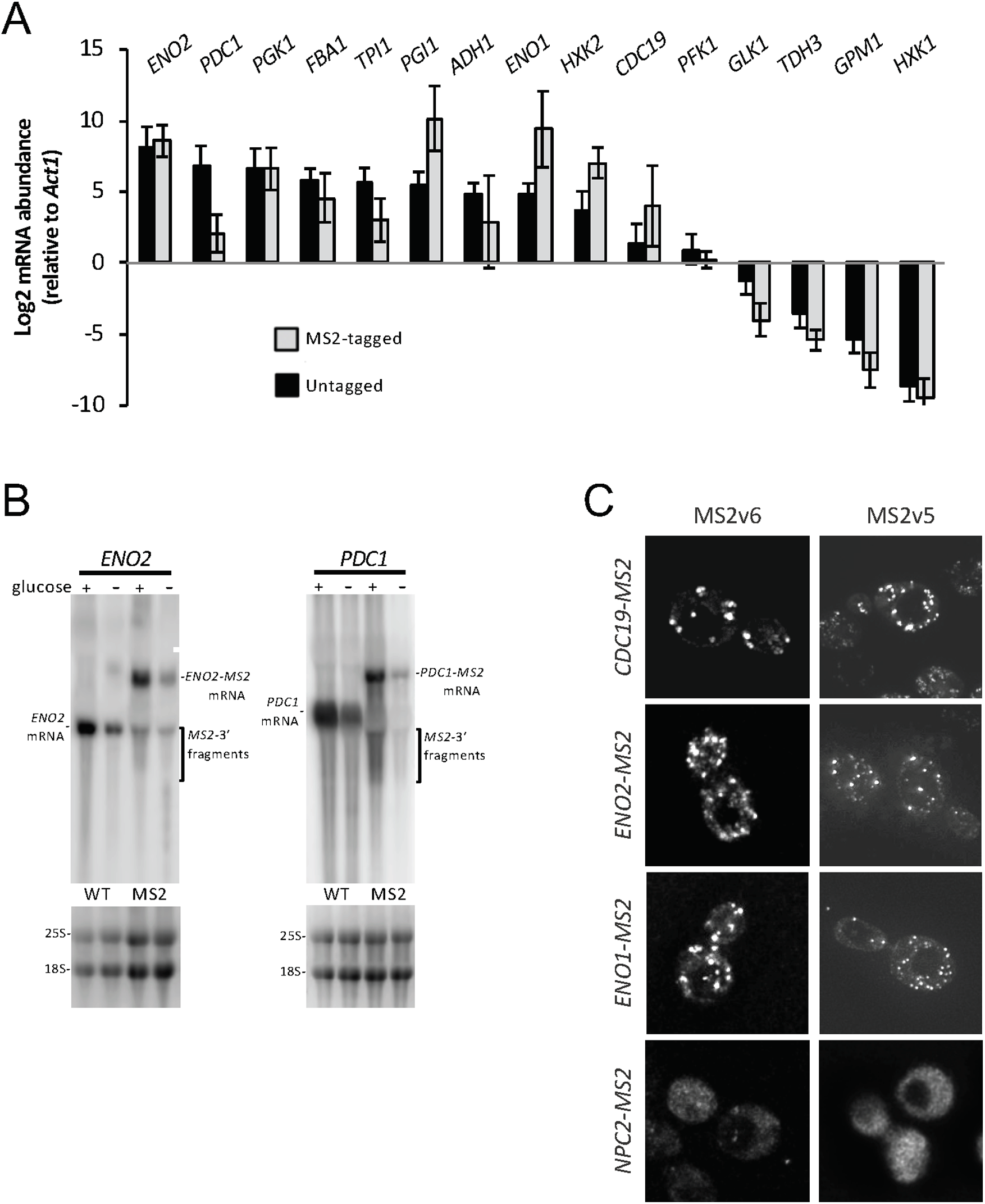
mRNA levels and the use of lower affinity MS2 stem loops support the premise that the glycolytic mRNAs are localized to RNA granules. (A) Graph representing the relative fold change of *MS2*-tagged mRNAs relative to untagged mRNA levels. Error bars represent ± SD. (B) Northern blots of *ENO2* and *PDC1* mRNA in glucose replete and starved conditions in untagged strains or strains bearing the *MS2* tag. (C) *z*-stacked epifluorescent images of *CDC19, ENO1* and *ENO2* mRNAs tagged with both the MS2v6 and MS2v5 stem loop systems co-expressing the relevant MS2-CP-GFP fusion. MS2v5 images are the same as those shown in Figure 1. *ENO1* and *CDC19* MS2v6 constructs contain 24x stem loops, *ENO2* contains 12x stem loops. Scale bar: 2 µm

